# *Mycobacterium tuberculosis* canonical virulence factors interfere with a late component of the TLR2 response

**DOI:** 10.1101/2021.10.06.463251

**Authors:** Amelia E. Hinman, Charul Jani, Stephanie C. Pringle, Wei R. Zhang, Neharika Jain, Amanda J. Martinot, Amy K. Barczak

## Abstract

For many intracellular pathogens, the phagosome is the site of events and interactions that shape infection outcome. Phagosomal membrane damage, in particular, is proposed to benefit invading pathogens. To define the innate immune consequences of this damage, we profiled macrophage transcriptional responses to wild-type *Mycobacterium tuberculosis* (Mtb) and mutants that fail to damage the phagosomal membrane. We identified a set of genes with enhanced expression in response to the mutants. These genes represented a late component of the TLR2-dependent transcriptional response to Mtb, distinct from an earlier component that included TNF. Expression of the later component was inherent to TLR2 activation, dependent upon endosomal uptake, and enhanced by phagosome acidification. Canonical Mtb virulence factors that contribute to phagosomal membrane damage blunted phagosome acidification and undermined the endosome-specific response. Profiling cell survival and bacterial growth in macrophages demonstrated that the attenuation of these mutants is partially dependent upon TLR2. Further, TLR2 contributed to the attenuated phenotype of one of these mutants in a murine model of infection. These results demonstrate two distinct components of the TLR2 response and identify a component dependent upon endosomal uptake as a point where pathogenic bacteria interfere with the generation of effective inflammation. This interference promotes TB pathogenesis in both macrophage and murine infection models.

## Introduction

Innate immune recognition of invading pathogens, typically driven by the interaction of pattern recognition receptors (PRRs) and pathogen-associated molecular patterns (PAMPs), requires recognition of microbial products at multiple subcellular sites. While some PRRs recognize PAMPs at a single site within the cell, other PRRs have the potential to bind PAMPs and initiate signaling from multiple sites. The mechanisms through which one PRR can recognize and respond distinctly to PAMPs at different subcellular sites is best understood for TLR4/LPS interactions [1-5]. Although principles elucidated with LPS and TLR4 are broadly thought to hold for other PAMP/PRR interactions, to date we have less insight into subcellular sites of signaling by other PRRs, the contribution of compartment-specific signaling in the response to complex microorganisms, and the pathogenic strategies employed to evade such compartmentalized signaling events.

TLR2, a receptor for bacterial cell wall lipoproteins, has been suggested to signal from the plasma membrane and endosomes, similar to TLR4. Endosome-specific TLR2 signaling in response to pathogenic bacteria has been partially explored using the model of *Staphylococcus aureus* taken up into macrophages [6]; in that work, TNF release was shown to be partially dependent upon TLR2 and dependent upon endosomal uptake. TLR2 activation has also been described to induce a type I interferon (IFN) transcriptional response from endosomes [7-10]. Overall, the mechanisms and physiological contexts in which compartment-specific TLR2 signaling occurs are unclear. In particular, it is unknown whether findings with *S. aureus* extend to other infectious agents and whether pathogens use strategies to prevent TLR2 signaling from the plasma membrane or endosomes.

*Mycobacterium tuberculosis* (Mtb) represents a model to study potential mechanisms of innate immune evasion, as this pathogen co-evolved with mammals and encodes multiple strategies of host manipulation. Mtb has a complex repertoire of PAMPs, and infection with Mtb is recognized by both membrane-bound and cytosolic PRRs. The specific complement of PRRs that drive the macrophage response to the intact bacterium and the subcellular sites of recognition of those PAMPs have not been clearly defined. The canonical Mtb virulence factors phthiocerol dimycocerosate (PDIM) and ESX-1 contribute to disruption of the macrophage phagosomal membrane upon infection [11-15]; we sought to leverage this shared pathogenic effect to gain insight into compartment-specific signaling in the macrophage response to Mtb.

To probe the relationship between Mtb-mediated phagosomal membrane damage and innate immune recognition of infection, we serially profiled the macrophage response to wild-type Mtb or PDIM and ESX-1 Mtb mutants, which fail to damage the phagosomal membrane. We found that the mutants elicited markedly enhanced expression of a cluster of inflammatory genes induced late after infection; expression was strictly dependent upon MYD88 and TLR2. TNF expression and release are commonly used as a marker of TLR activation; however, we found that induction of TNF occurred with an earlier set of TLR2-dependent genes and differed minimally between the response to wild-type Mtb and our mutants. We thus hypothesized that infection with Mtb elicits a two-component TLR2-dependent response, and that the later component of the response is preferentially blunted by Mtb factors that damage the phagosomal membrane. Treatment of macrophages with synthetic TLR2 ligand elicited a similar two-component transcriptional response, suggesting that these components are fundamental facets of TLR2 signaling rather than pathogen-specific. Induction of the early component of the TLR2 response was similar in the presence of endosomal uptake inhibitors; in contrast, the later component was markedly diminished by inhibition of endosomal uptake. Induction of the endosome-specific response was dependent upon phagosome acidification. We found that Mtb factors known to damage the phagosomal membrane contributed to Mtb-induced limitation of phagosome acidification, which in turn limited production of the late component of the TLR2 response. Consistent with published reports, PDIM-mutant and ESX-1-mutant Mtb had attenuated virulence phenotypes in wild-type macrophages, with reduced macrophage cytotoxicity and reduced bacterial growth. Both of these attenuated phenotypes were partially reversed in TLR2-/-macrophages, suggesting that TLR2-dependent responses contribute to the attenuation of these mutants. As expected, PDIM-mutant Mtb had attenuated infection phenotypes in C57BL/6J mice, with reduced bacterial growth and lung infiltrates. In contrast, in TLR2-/-mice PDIM-mutant Mtb grew more robustly and caused pulmonary infiltrates more similar to wild-type Mtb. These results support a model in which PDIM and ESX-1 contribute to Mtb virulence in part by blunting a protective endosome-specific component of the TLR2-dependent response to infection.

## Results

### Macrophage infection with Mtb PDIM or ESX-1 mutants elicits enhanced expression of an inflammatory transcriptional program

The Mtb ESX-1 protein secretion system has long been known to mediate phagosomal membrane damage [13, 15]; the mycobacterial lipid phthiocerol dimycocerosate (PDIM) was more recently found to play a similar role in infection [11, 12, 14]. To understand how the capacity to damage the phagosomal membrane shapes the innate immune response to Mtb, we sought to leverage the similar effects of PDIM and ESX-1-mediated secretion within the macrophage. We thus compared the response to wild-type Mtb with the response to PDIM and ESX-1 mutants, with the goal of identifying facets of the macrophage response to Mtb impacted by phagosomal membrane damage. To enable meaningful comparison, we first tested whether wild-type Mtb, PDIM mutants, and ESX-1 mutants were taken up similarly into bone marrow-derived macrophages (BMDM). While the possibility of PDIM facilitating uptake into macrophages has been raised [16, 17], using either CFU or flow cytometry we found that wild-type Mtb strain H37Rv, PDIM mutants, and ESX-1 mutants were taken up at similar rates (**Supp. Fig. S1A-B**).

To define the macrophage transcriptional programs induced by Mtb, we infected BMDM with wild-type Mtb and performed comprehensive transcriptional profiling at 4, 8, 12, and 16 hours post-infection (**Supp. Table 1**). Focusing on the 907 genes changed two-fold upon infection, genes could be categorized into three clusters with distinct patterns of expression (**Fig. 1A**). Cluster 1 genes were progressively induced, and cluster 2 genes were progressively repressed over time after infection. Cluster 3 genes were induced upon infection and peaked at 8-12 hours before waning. To identify the facets of the macrophage response most impacted by phagosomal membrane damage, we next compared this baseline response to wild-type Mtb with the response to PDIM or ESX-1 mutants (**Supp. Table 1**). For this comparison, we used PDIM mutants we had demonstrated in previous work to lack PDIM production (Δmas and ΔppsD) and an ESX-1 mutant we had demonstrated lacked ESX-1 secretion but produced and properly localized PDIM (Tn::eccCa1) [12]. Comparing the macrophage response to our mutants with the response to wild-type Mtb, we found that expression of genes in clusters 2 and 3 was similar in response to each of the three strains. In contrast, expression of genes in cluster 1 was markedly impacted by loss of PDIM or ESX-1. However, not all genes in the cluster responded similarly to the mutants; classifying genes in this cluster based on their response to PDIM and ESX-1 mutants clearly distinguished two subclusters (**Fig. 1B**). Induction of genes in subcluster 1A was markedly diminished in response to PDIM and ESX-1 mutants relative to wild-type Mtb (**Fig. 1B**). Ingenuity Pathway Analysis [18] of genes in this subcluster predicted STAT1, IRF3, interferon-α, and IFNAR as upstream regulators with high confidence, suggesting that these genes comprised the type I IFN response. Manual inspection confirmed that interferon stimulated genes were highly represented in subcluster 1A. These results were consistent with previous work demonstrating that the macrophage type I IFN response to Mtb is dependent upon ESX-1-mediated secretion [19] and PDIM [12].

**Figure 1.**
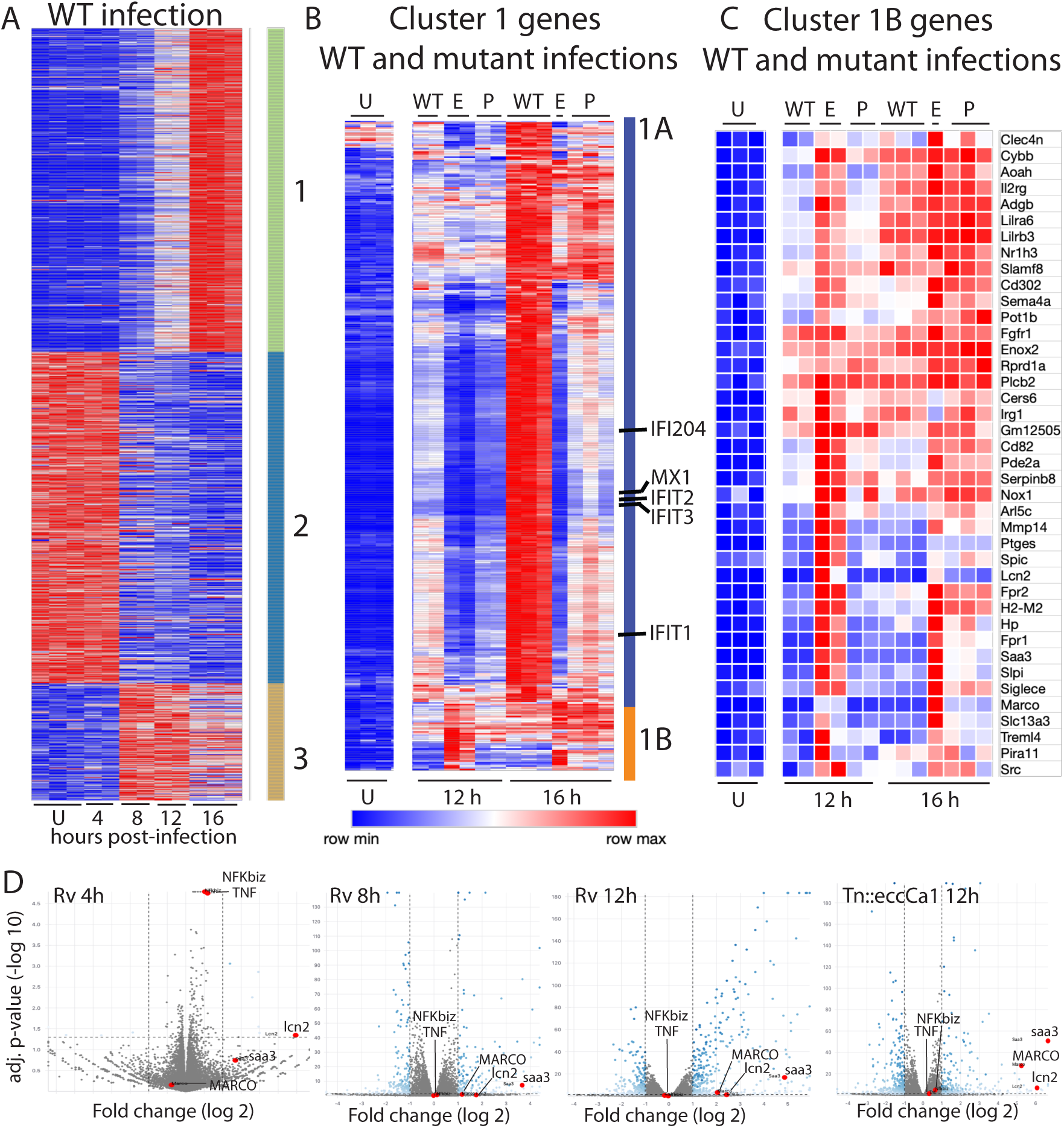
Macrophage infection with Mtb PDIM or ESX-1 mutants reveals two subclusters of genes differentially expressed relative to infection with wild-type Mtb. (A-C) C57BL/6J BMDM were infected with wild-type Mtb H37Rv (“WT”), the ESX-1 core complex mutant *Tn::eccCa1* (“E”), or the PDIM production mutant *Δmas* (“P”) at an MOI of 2:1. At 4, 8, 12, and 16 hours post-infection, RNA was harvested for RNAseq. Sequencing libraries not passing QC metrics were excluded from further analysis. (A) Genes were clustered based on similarity of expression in response to WT Mtb. (B, C) Cluster 1 genes from *A* were subclustered based on the response to WT Mtb and the mutants. Uninfected, 12h, and 16h timepoints shown. (A-C) Blue-red gradient reflects relative expression within each row. (D) Volcano plots for the indicated conditions. TNF and co-regulated gene NFKbiz and subcluster 1B genes saa3, MARCO, lcn2 are indicated on each graph. RNAseq experiment performed once.

In contrast to subcluster 1A, expression of genes in subcluster 1B was enhanced in response to PDIM or ESX-1 mutants relative to wild-type Mtb at later timepoints (**Fig. 1C-D, Fig. 2A**). Strikingly, this subcluster included multiple genes important for the host response to Mtb, including MARCO [20], prostaglandin E synthase [21-24], lipocalin 2 [25, 26], Irg1 [27, 28] and matrix metalloproteinase 14 [29]. The kinetics of overall induction and enhanced expression observed in response to the PDIM and ESX-1 mutants were independent of MOI, as the same patterns were observed infecting with MOI 2:1 and 10:1 (**Figs. 2A-B**). Enhanced expression was similarly elicited in BALB/c BMDM, suggesting that the enhanced response is independent of the background genetic inflammatory state of the cells (**Supp. Fig. S1C**). Enhanced expression was observed regardless of the specific PDIM or ESX-1 mutation, including ESX-1 core complex mutant *Tn::eccCa1*, secreted effector mutants (*ΔesxB* and *Tn::espC*), mutants in the synthetic pathway for distinct components of PDIM (*Δmas* and *ΔppsD*), and a PDIM transport mutant (*Tn::drrC*) (**Fig. 2C**). Complementation of the disrupted ESX-1 or PDIM gene restored expression to wild-type levels (**Fig. 2D**). PDIM and ESX-1 mutants are both known to be attenuated for growth in macrophages [30, 31]; we thus considered the possibility that any attenuated mutant would elicit enhanced expression of subcluster 1B genes. To test this possibility, we obtained a well-characterized Mtb mutant. The deleted gene, Δ*pckA*, catalyzes the first step in gluconeogenesis, and the mutant is highly attenuated for growth in macrophages [32] (**Supp. Fig. S1D**). Infection with the Δ*pckA* mutant did not elicit enhanced expression of subcluster 1B genes (**Supp. Fig. S1E**), demonstrating that the enhanced response to PDIM or ESX-1 mutants is not a general response to mutants with impaired intracellular survival. These results suggest that PDIM and ESX-1 functions blunt induction of an inflammatory transcriptional program that includes multiple genes individually linked to control of TB infection.

**Figure 2.**
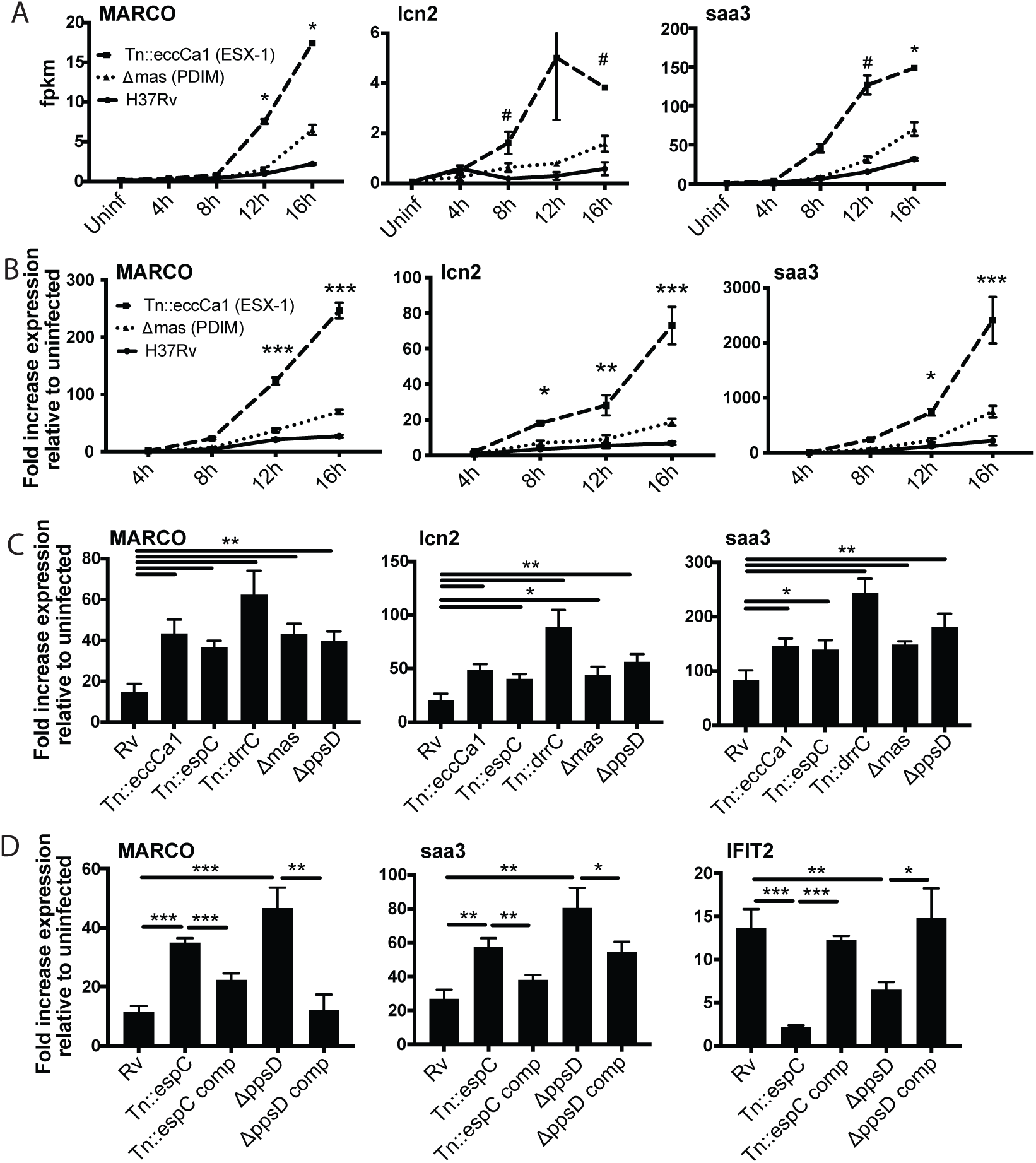
Infection with Mtb PDIM or ESX-1 mutants elicits enhanced expression of an inflammatory transcriptional program. (A) fpkm from RNAseq data (MOI 2:1) for representative genes from subcluster 1B. * p-value < 0.01 or # p-value < 0.05 for the comparison of PDIM (16h) or ESX-1 (12h) mutant-infected with Rv-infected, unpaired two-tailed t-test. (B) C57BL/6J BMDM were infected with the indicated strains at an MOI of 10:1. RNA was harvested at the indicated timepoints, and expression of the indicated genes was profiled by qPCR relative to GAPDH control. (C-D) C57BL/6J BMDM were infected with the indicated Mtb strains at an MOI of 2:1. RNA was harvested 24 hours post-infection, and expression of the indicated genes relative to GAPDH control was profiled using qPCR. (B-D) Mean +/-SD of 4 replicates. *p-value < 0.01, **p-value < 0.001, ***p-value < 0.0001 unpaired two-tailed t-test. RNAseq experiment (A) performed once, (B) one of two independent experiments (C-D) one of three independent experiments.

### PDIM and ESX-1 blunt the later component of a biphasic TLR2-dependent transcriptional response to Mtb

Ingenuity Pathway Analysis of genes in subcluster 1B predicted MYD88 and NF-kB as upstream regulators. To test these predictions and define upstream regulators, we profiled expression in BMDM from knockout mice. Consistent with pathway predictions, expression in response to either wild-type Mtb or the mutants was lost in MYD88-/-BMDM (**Fig. 3A**). MYD88 functions as a signaling adapter for TLRs; we next sought to identify the relevant upstream TLR. Mtb produces multiple potential TLR2 antigens [33-39]; we thus tested a role for TLR2 as the relevant upstream TLR. Similar to findings for MYD88, expression of representative genes upon infection with wild-type Mtb or PDIM or ESX-1 mutants was entirely lost in TLR2-/-BMDM (**Fig. 3A**). In contrast, TLR4-/-knockout BMDM responded similarly to wild-type BMDM **(Fig. 3B)**. These results confirmed MYD88 and TLR2 as upstream regulators of macrophage transcriptional response component blunted by PDIM and ESX-1 function.

**Figure 3.**
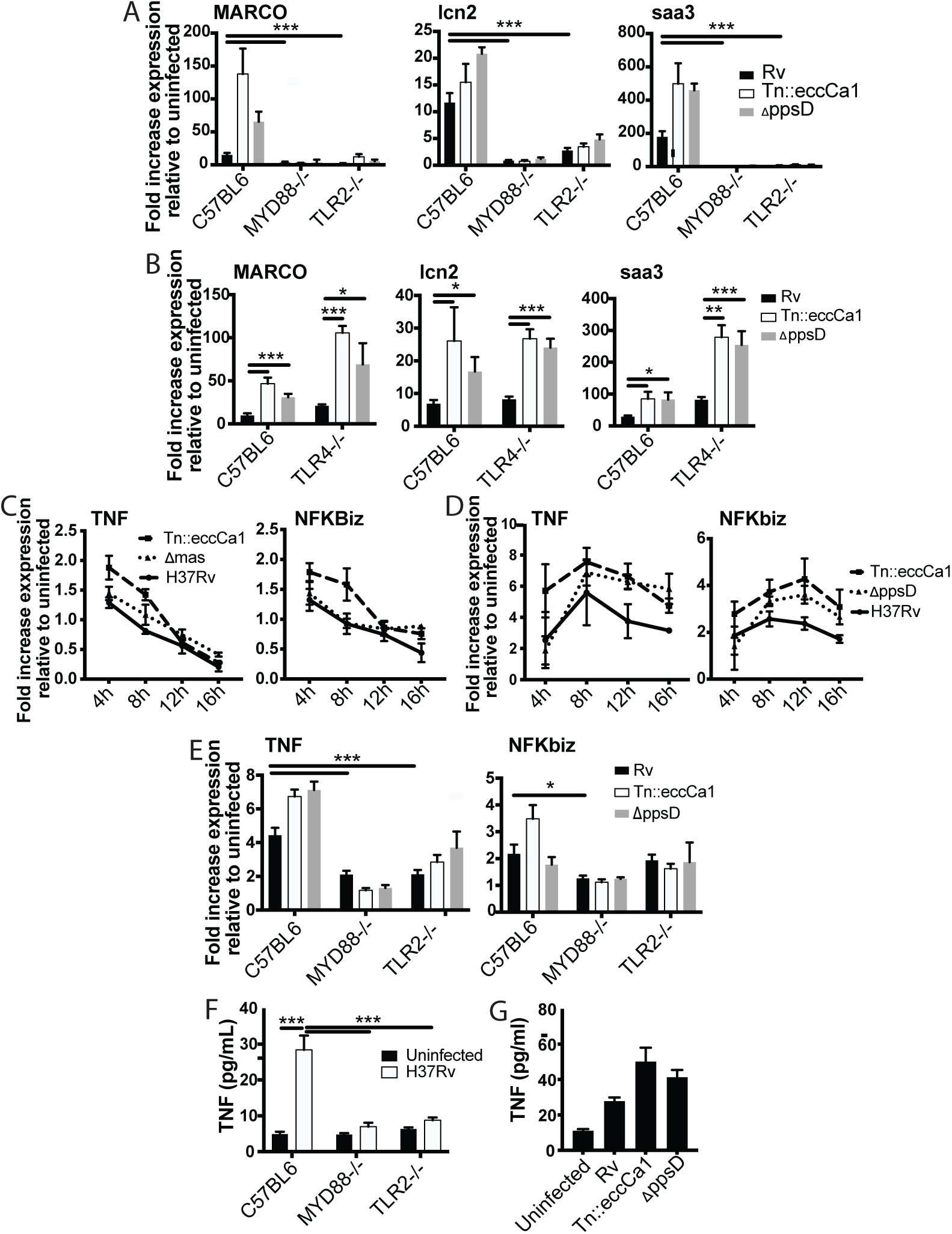
The identified two-component inflammatory response to Mtb is dependent upon MYD88 and TLR2. (A-B) The indicated BMDM were infected with the indicated Mtb strains at an MOI of 2:1. RNA was harvested 24 hours post-infection. (C-D) C57BL/6J BMDM were infected with the indicated Mtb strains at an MOI of 2:1 (C) or 10:1 (D). RNA was harvested at the indicated timepoints post-infection. (E) The indicated BMDM were infected with the indicated Mtb strains at an MOI of 5:1. RNA was harvested 6 hours after infection. (F-G) The indicated (F) or C57BL/6J (G) BMDM were infected with the indicated Mtb strains at an MOI of 5:1. Supernatants were harvested 24 hours post-infection, and TNF was quantified by ELISA. Mean +/-SD of 4 replicates. *p-value < 0.01, ***p-value < 0.0001 unpaired two-tailed t-test. (A, C, E) one of three independent experiments (B, D, F-G) one of two independent experiments.

Given the preponderance of work using TNF as a marker of TLR activation, we next examined the relationship between TLR2, TNF expression, and PDIM and ESX-1 in the macrophage response to Mtb. TNF did not cluster with 1B genes; instead TNF was in cluster 3 (**Fig. 1A**) together with genes minimally impacted by PDIM and ESX-1. Our RNAseq data additionally demonstrated that expression of TNF peaked earlier post-infection than expression of subcluster 1B genes and then waned. At an MOI of 2:1, we found that induction of TNF was very modest (less than two-fold at the time of peak induction, **Fig. 3C**), limiting our ability to reliably any decrease in TNF expression. Infection at an MOI of 10:1 modestly enhanced expression of TNF and delayed the time of peak expression (**Fig. 3D**). At both MOI 2:1 and 10:1, we found that TNF expression was very modestly impacted by PDIM and ESX-1 (**Fig. 3C-D**). MOI of 5:1 gave similar kinetics and magnitude of TNF expression to MOI of 10:1 (**Supp. Fig. S2A**); we thus selected an MOI of 5:1 to minimize macrophage cell death while allowing us to reliably measure any impact of experimental interventions on TNF expression. TNF expression was partially lost in TLR2 and MYD88 knockout BMDM (**Fig. 3E**), suggesting that additional PRRs likely contribute to TNF expression in the response to Mtb. We hypothesized that TNF was part of a broader early TLR2-dependent transcriptional program; clustering genes across all infection conditions identified 17 genes with expression highly correlated with TNF (**Supp. Fig. S2B**). Testing expression of an additional representative gene from that cluster, NFKbiz, confirmed a pattern of expression similar to TN**F**. Similar to TNF, expression was minimally impacted by PDIM and ESX-1 and was partially dependent upon MYD88 and TLR2 (**Fig. 3C-E**). For Mtb infection of macrophages, TNF transcription and release have been described as potentially dissociated [40]. However, we found that similar to TNF transcription, TNF release upon Mtb infection was partially dependent upon MYD88 and TLR2 (**Fig. 3F**) and very modestly increased upon infection with PDIM- or ESX-1-mutant Mtb (**Fig. 3G**). Our results thus suggest that Mtb infection of macrophages induces distinct early and late TLR2-dependent transcriptional responses, and that canonical Mtb virulence factors that interfere with phagosomal membrane integrity preferentially blunt the later response.

### The observed biphasic transcriptional response is a fundamental feature of TLR2 signaling

We reasoned that the two-component TLR2 response we observed upon Mtb infection either could be pathogen-specific or could reflect a fundamental feature of TLR2 signaling. To distinguish between these possibilities, we treated BMDM with PAM3CSK4 or PAM2CSK4, synthetic agonists of TLR1/TLR2 and TLR2/TLR6, respectively, and profiled expression of genes representative of the early and late TLR2-dependent response to Mtb. Treatment with either synthetic ligand elicited expression of genes in the early component of the TLR2-dependent response to Mtb with a similar pattern of expression, peaking at 2-4 hours post-treatment before waning (**Fig. 4A, Supp. Fig. S3A)**. PAM3CSK4 or PAM2CSK4 also elicited expression of genes in the later TLR2-dependent response component with delayed kinetics, evident by 4 hours post-infection but continuing to increase through 24 hours (**Fig. 4B, Supp. Fig. S3B**) As with Mtb infection, synthetic ligand-dependent expression of genes representative of both the early and late clusters was dependent upon TLR2 and the signaling adapter MYD88 (**Fig. 4C-D**). PIM6 has been described to be the most potent TLR2 agonist in the Mtb PAMP repertoire [41]; we found that treating BMDM with purified PIM6 similarly induced the early and late response genes in a TLR2-dependent manner (**Fig. 4E-F**). The two observed components of the TLR2-dependent response to Mtb infection thus appear to reflect inherent dynamics of TLR2 signaling in macrophages rather than dynamics specific to recognition of intact Mtb. None of the known adapters TRIF, TRAM, or TIRAP were required for induction of the late response (**Supp. Fig. S3C-D**).

**Figure 4.**
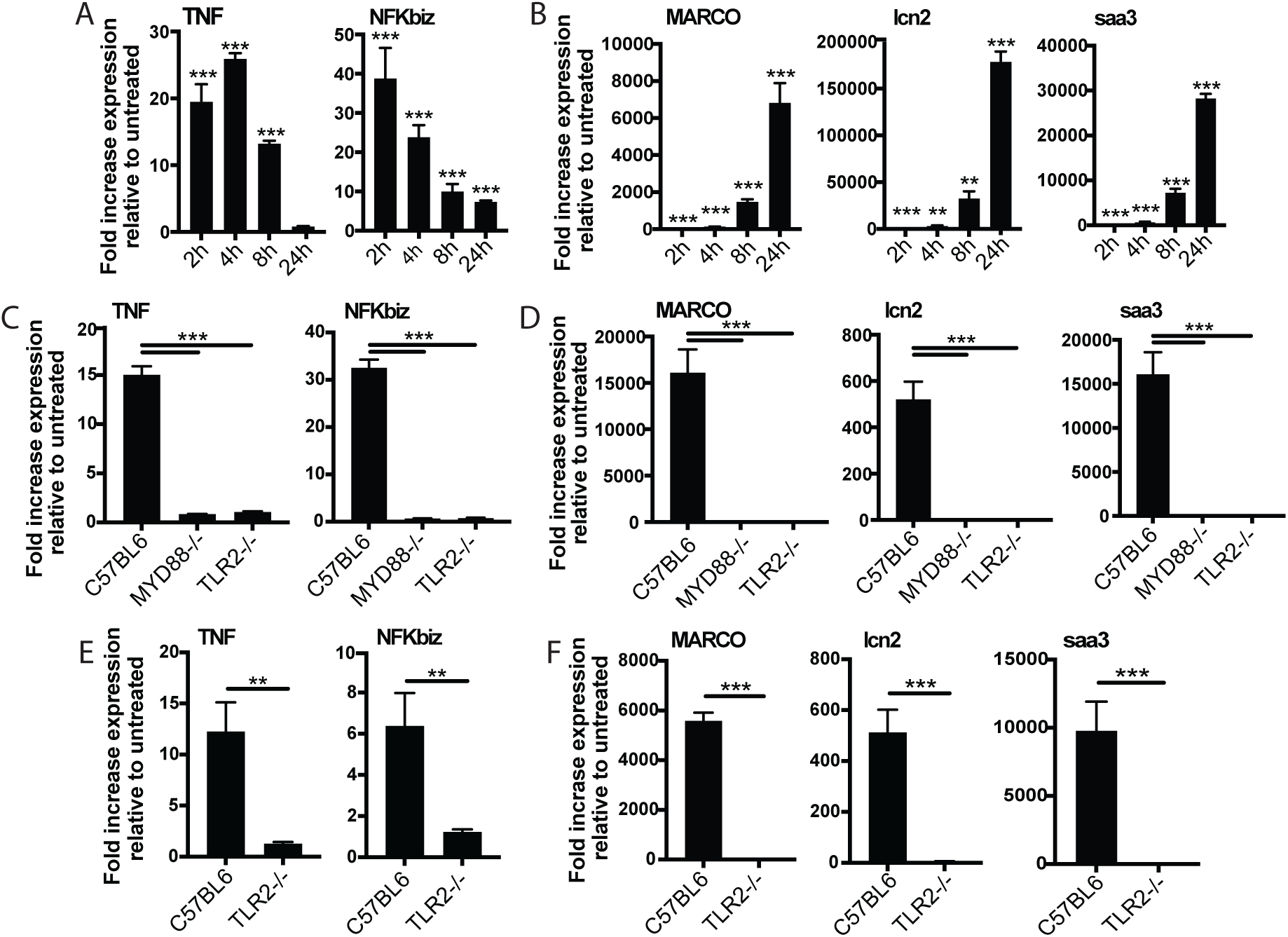
The two-component response is a fundamental feature of TLR2 activation. C57BL/6J (A-B) or the indicated (C-F) BMDM were treated with PAM3CSK 1μg/ml (A-D) or PIM6 1μg/ml (E-F) and RNA was harvested at the indicated timepoints (A-B), 2 hours (C, E), or 24 hours (D, F). qPCR was performed to quantitate expression of the indicated genes relative to GAPDH control. Mean +/-SD for 4 replicates. **p-value < 0.001, ***p-value < 0.0001 unpaired two-tailed t-test. (A-B) one of two independent experiments, (C-F) one of three independent experiments.

### The later component of the TLR2-dependent transcriptional response requires endosomal uptake

We next considered possible determinants of the two distinct TLR2 response components. We first considered that expression of genes in the later component may be driven by signaling through the TNF receptor initiated by the early component; however, late response component genes were expressed similarly in wild-type and TNF receptor knockout BMDM (**Supp. Fig. S4A**). We then considered alternate hypotheses. PDIM and ESX-1 function preferentially undermine the second component of the response, and both interact with the phagosomal membrane. We thus hypothesized that the determinant of the distinct components may be spatial, with expression of the two sets of genes initiated at distinct subcellular sites. To distinguish between surface-initiated signal and endosome-specific signal, we used the dynamin inhibitor dynasore [42]. Dynamin is required for the final step of formation of the endocytic vesicle, and in the context of LPS recognition by TLR4, dynamin has been used to dissect compartment-specific aspects of signaling [4, 43, 44]. In the context of TLR2 on macrophages, dynamin does not change the surface and endosomal distribution of the receptor, but blocks uptake of PAM3CSK4 into the endosome [45]. We thus used dynasore and PAM3CSK4 to test whether uptake of synthetic TLR2 ligand into the endosome is required for activation of either component of the response. PAM3CSK4-induced expression of genes in the early cluster was not significantly changed by dynasore pre-treatment (**Fig. 5A**). In contrast, PAM3CSK4-induced expression of genes in the late cluster was markedly reduced (**Fig. 5B**).

**Figure 5.**
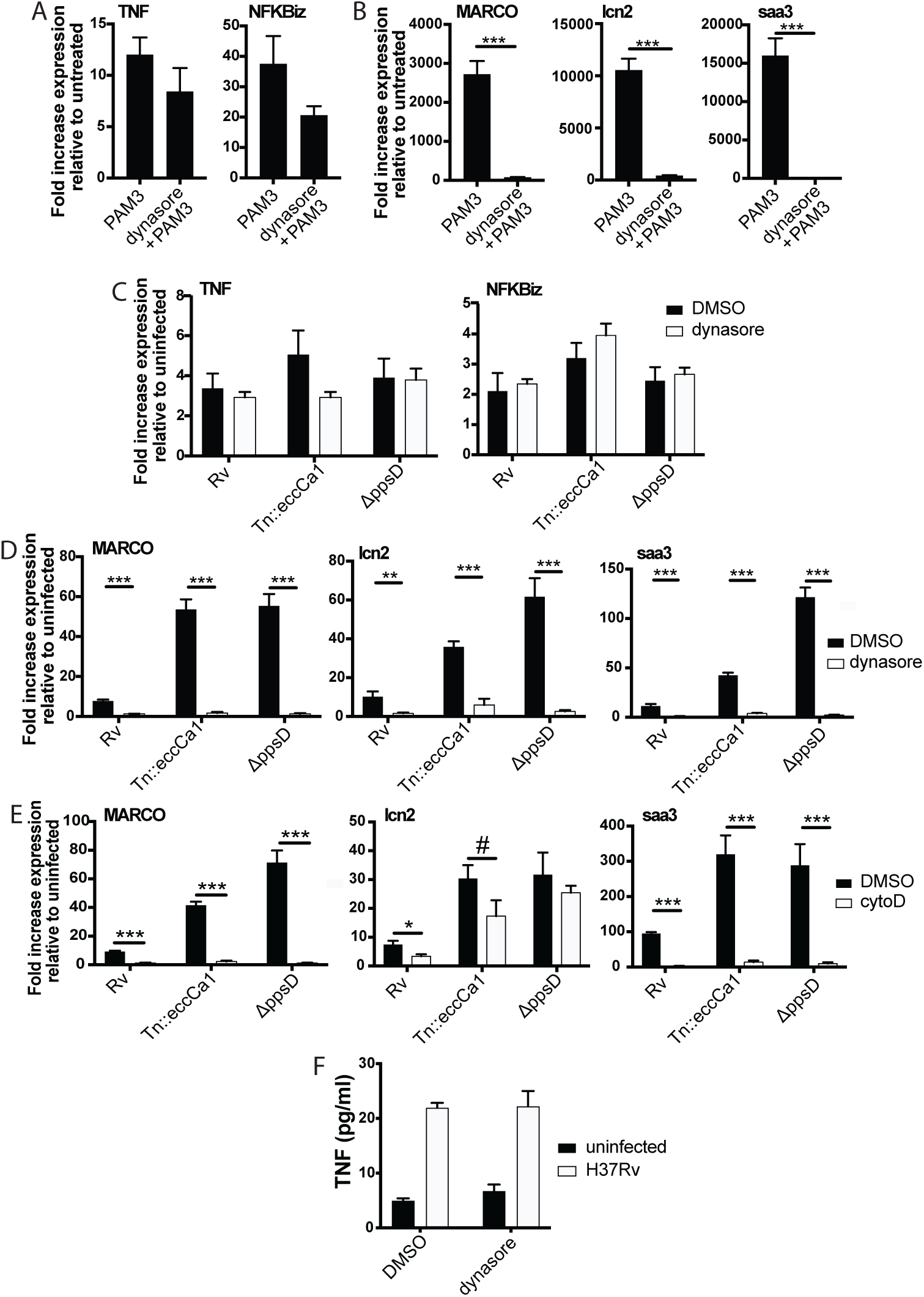
The later component of the TLR2-dependent transcriptional response requires endosomal uptake. Where indicated, C57BL/6J BMDM were pre-treated with dynasore 80μM or cytochalasin D 10μM. Cells were then treated with PAM3CSK4 1ug/ml (A-B) or infected with the indicated Mtb strains at an MOI of 5:1 (C) or 2:1 (D-E). RNA was harvested at 2 hours (A) 6 hours (C) or 24 hours (B,D-E). qPCR was performed to quantitate expression of the indicated genes relative to GAPDH control. (E, F) BMDM were infected with the indicated Mtb strains at an MOI of 5:1. Supernatants were harvested 24 hours post-infection, and TNF was quantified by ELISA. Mean +/-SD for 4 replicates. #p-value < 0.05, *p-value < 0.01, ***p-value < 0.0001 unpaired two-tailed t-test. (A-E) one of three independent experiments, (F) one of two independent experiments.

We next wanted to test whether the compartment-specificity of the two components of the TLR2-dependent proinflammatory response to synthetic ligand is similar for the response to Mtb. The effect of dynasore on Mtb uptake has not previously been tested; using gentamicin protection assays, we found that dynasore significantly decreased Mtb uptake **(Supp. Fig. S4B-C)**. We then tested the effect of inhibiting Mtb uptake on expression of genes in the early and late clusters. Similar to treatment with synthetic ligand, inhibition of Mtb uptake with dynasore left expression of genes in the early TLR2-dependent cluster largely preserved (**Fig. 5C**), but significantly blunted expression of genes in the late cluster (**Fig. 5D**). Results were similar when BMDM were pre-treated with the actin polymerization inhibitor cytochalasin D (**Fig. 5E, Supp. Fig. S4C**), which has previously been used to distinguish innate immune signaling pathways initiated from the cell surface vs. the endosome [6, 7, 9]. Similar to the patterns observed for TNF transcription, dynasore pre-treatment had minimal impact on TNF release (**Fig. 5F**). These results suggest that while expression of early cluster genes can be initiated from the plasma membrane, expression of genes in the later cluster is dependent upon endosomal uptake.

Induction of type I IFNs in response to TLR2 activation has been previously linked to endosomal uptake [7-10]. We found that stimulation of BMDM with TLR2 agonists modestly induced type I IFNs (**Supp. Fig. S5A-B**); this induction was dependent upon TLR2 (**Supp. Fig. S5C**) and partially inhibited by dynasore pre-treatment (**Supp Fig. S5D**). However, the kinetics and magnitude of induction of IFN-β were distinct from the late pro-inflammatory component of the TLR2 response, suggesting that the endosome-specific pro-inflammatory response is distinct from the type I IFN response. Consistent with established models, induction of the type I IFN response to Mtb was independent of TLR2 (**Supp. Fig. S5E**). Our results suggest that while induction of the second component of the TLR2-dependent response is similar between Mtb and purified or synthetic TLR2 ligand, induction of type I IFNs is not part of this shared response.

### Full activation of the endosome-specific TLR2 response is dependent upon phagosome acidification

We next sought to understand how PDIM and ESX-1 function might undermine induction of the second component of the TLR2 response. Both PDIM and ESX-1 are required for induction of the type I IFN response to Mtb [12, 19] (**Supp. Fig. 5F**), and interference between induction of type I IFNs and NF-ΚB at the transcription factor level has previously been proposed in the macrophage response to other pathogens [46]. We thus hypothesized that PDIM and ESX-1-facilitate induction of type I IFNs, and that type I IFN-activated transcription factors interfere with binding of NF-ΚB-dependent transcription factors that contribute to the later component of the TLR2 response. To test this hypothesis, we profiled the macrophage response to Mtb in macrophages unable to mount a type I IFN response to infection. STING is strictly required for the type I IFN response to Mtb upstream of IRF3 activation [13] (**Supp. Fig. S5E**). We predicted that if type I IFN-activated transcription factors blunt the TLR2 response, the response to wild-type Mtb would be increased in STING knockout BMDM relative to wild-type BMDM. We additionally predicted that the response to wild-type Mtb and PDIM or ESX-1 knockouts would be equivalent in STING knockout BMDM, as the type I IFN response would be similarly absent in response to all three Mtb strains. In fact, neither prediction tested correct (**Supp. Fig S5G**), suggesting that the mechanism through which PDIM and ESX-1 blunt the TLR2-dependent response to Mtb is independent of their role in type I IFN induction.

We then considered other ways that PDIM and ESX-1 function might interfere with the TLR2 response. Both PDIM and ESX-1 are required for phagosomal membrane damage [11, 13, 14]. Candida-mediated phagosomal membrane damage and sterile phagosomal membrane damage have both been described to interfere with phagosome acidification, potentially because of loss of the proton gradient across the membrane at sites of damage [47, 48]. We reasoned that PDIM- and ESX-1-mediated membrane damage might similarly contribute to the known limitation of acidification in Mtb-containing phagosomes. An ESX-1 mutant in *M. marinum* has in fact previously been shown to reside in a more highly acidified macrophage phagosome than wild-type *M. marinum* [49]. Providing suggestive evidence for a link between phagosome acidification and the TLR2 response, inhibitors of phagosome acidification limit the MYD88-dependent response to *Staphylococcus aureus* [6]; this effect was attributed to a requirement for cathepsin activation within acidified lysosomes to process intact *S. aureus* and release TLR agonists. We thus hypothesized that PDIM and ESX-1 mediated membrane damage contributes to the limitation of phagosome acidification, and that limitation then impacts endosome-specific TLR2 activation.

To first test whether PDIM and ESX-1 function impact phagosome pH, we used the pH-sensitive fluorescent dye pHrodo. pHrodo labeling of Mtb has previously been used to quantify phagosomal pH around the mycobacterium [50]. We labeled PDIM-mutant, ESX-1-mutant, or wild-type Mtb expressing GFP with pHrodo, then infected BMDM. CellProfiler [51] image analysis was used to identify GFP-Mtb; the corresponding pHrodo mean fluorescent intensity for each identified bacterium was then measured (**Supp. Fig. S6A-B**). At 6, 12, or 24 hours post-infection, pHrodo mean fluorescent intensity around PDIM or ESX-1 mutant Mtb was significantly higher than around wild-type Mtb (**Fig. 6A**), suggesting that phagosomes containing PDIM- or ESX-1-mutant Mtb becomes relatively more acidic than phagosomes containing wild-type Mtb.

**Figure 6.**
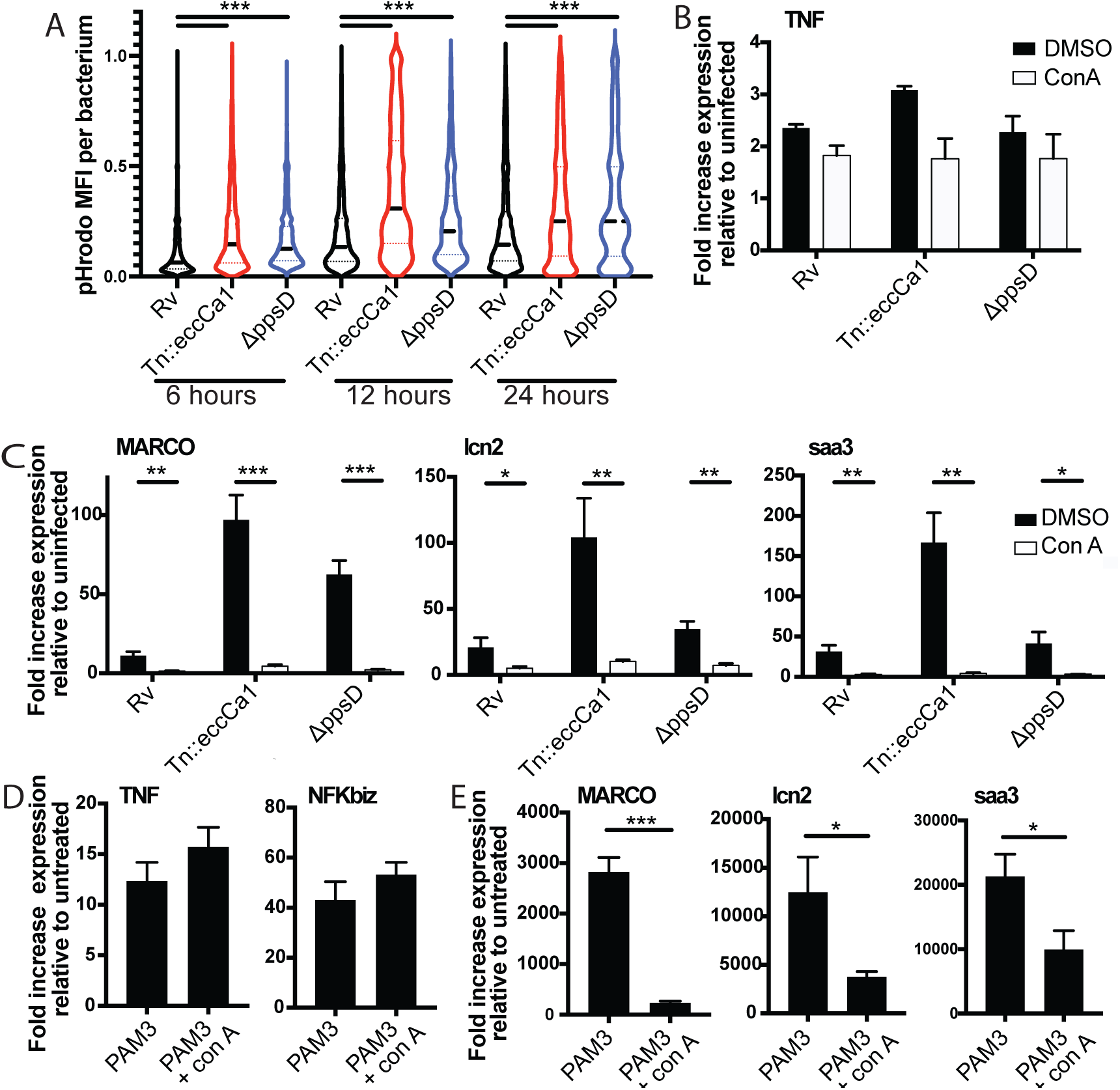
Full activation of the endosome-specific TLR2 response is dependent upon phagosome acidification. (A) The indicated Mtb strains expressing GFP were labeled with pHrodo and used to infect C57BL/6J BMDM at an MOI of 3:1. After 4 hours, cells were washed to remove extracellular bacteria. Cells were fixed at 6, 12, and 24 hours post-infection and imaged. Bacteria were identified based on GFP signal, and pHrodo mean fluorescence intensity was measured around each bacterium. A minimum of 1703 bacteria were analyzed per group. (B-C) C57BL/6J BMDM were pretreated with concanamycin A 50μM, then infected with the indicated Mtb strains at an MOI of 5:1 (B) or 2:1 (C). (C, D) C57BL/6J BMDM were pretreated with concanamycin A 50μM, then stimulated with PAM3CSK4 1μg/ml. RNA was harvested at 6 hours (B), 24 hours (C, E) or 2 hours (D) post-infection. qPCR was performed to quantitate expression of the indicated genes relative to GAPDH control. Mean +/-SD for 4 replicates. # p-value < 0.03, * p-value < 0.01, ** p-value < 0.001, ***p-value < 0.0001, unpaired two-tailed t-test. (A-E) one of three independent experiments.

We then tested whether phagosome acidification enhances endosomal TLR2 signaling. We first confirmed that pre-treatment with concanamycin A, which inhibits the vacuolar ATPase, limits phagosome acidification in BMDM using both zymosan beads (**Supp. Fig. S6C**) and infection with pHrodo-labeled PDIM and ESX-1 mutants (**Supp. Fig. S6D**). We then pre-treated BMDM with concanamycin A prior to infection with Mtb. We found that while expression of the early cluster of genes was not significantly changed (**Fig. 6B**), expression of the second cluster of genes was markedly diminished in the presence of concanamycin A (**Fig 6C**). These results suggested that phagosome acidification enhances the endosomal component of the TLR2-dependent response to Mtb. If acidification primarily drives the release of antigens from intact Mtb, as described for *S. aureus* [6], we would expect this acidification to be relevant upon infection with intact bacteria, but dispensable for the response to synthetic TLR2 ligand. Expression of genes in the early component of the response to synthetic TLR2 ligand was not changed by the addition of concanamycin A (**Fig. 6D**). However, expression of genes in the later component of the response was markedly diminished by the addition of concanamycin A (**Fig. 6E**). Similar results were obtained for the Mtb TLR2 ligand PIM6 (**Supp. Fig. S6E-F**). These results suggest that the late component of TLR2 signaling is dependent on phagosome acidification entirely independent of the capacity to process pathogen and release TLR2 agonists. Taken together, our results are consistent with the hypothesis that Mtb-mediated damage of the phagosome membrane blunts the endosome-specific TLR2 response by limiting phagosome acidification. Further, the dependence on phagosome acidification suggests a potential mechanism through which compartment-specific TLR2 signaling is regulated. Signaling of the endosome-restricted TLRs, TLR7 and TLR9, is in fact strictly dependent upon endosomal acid-activated proteases [52, 53], offering precedent for pH as a regulator of compartment-specific TLR signaling.

### PDIM and ESX-1 modulate TLR2-dependent infection outcomes in macrophages

We next sought to understand whether the interaction between PDIM/ESX-1 and TLR2 contributes to infection outcomes in macrophages. Mtb infection has been shown to drive macrophage cell death, including apoptosis [54], necrosis [55], and ferroptosis [56]. PDIM has been described to specifically contribute to macrophage necrosis [14]; ESX-1 has also been shown to contribute to macrophage cell death after infection [11, 57]. In one study, pre-treatment of macrophages with a TLR4 or TLR2 agonist reduced Mtb-induced cell death [58]. Consistent with previous reports, we found that in wild-type macrophages PDIM and ESX-1 mutants induced less cell death than wild-type Mtb or complemented mutants (**Fig. 7A-B**) We hypothesized that PDIM/ESX-1 interference with the late component of the TLR2 response might contribute to the cell death induced by wild-type Mtb; in that case, we would expect the enhanced macrophage survival observed upon infection with the PDIM or ESX-1 mutants to be lost or diminished in TLR2-/-macrophages. Infection with wild-type Mtb induced a similar degree of cell death in wild-type and TLR2-/-macrophages. However, the resistance to cell death observed in wild-type macrophages infected with PDIM or ESX-1 mutants was partially lost in TLR2-/-macrophages (**Fig. 7A-B**). These results suggest that PDIM and ESX-1 interference with TLR2-dependent responses contributes to macrophage cell death following infection.

**Figure 7.**
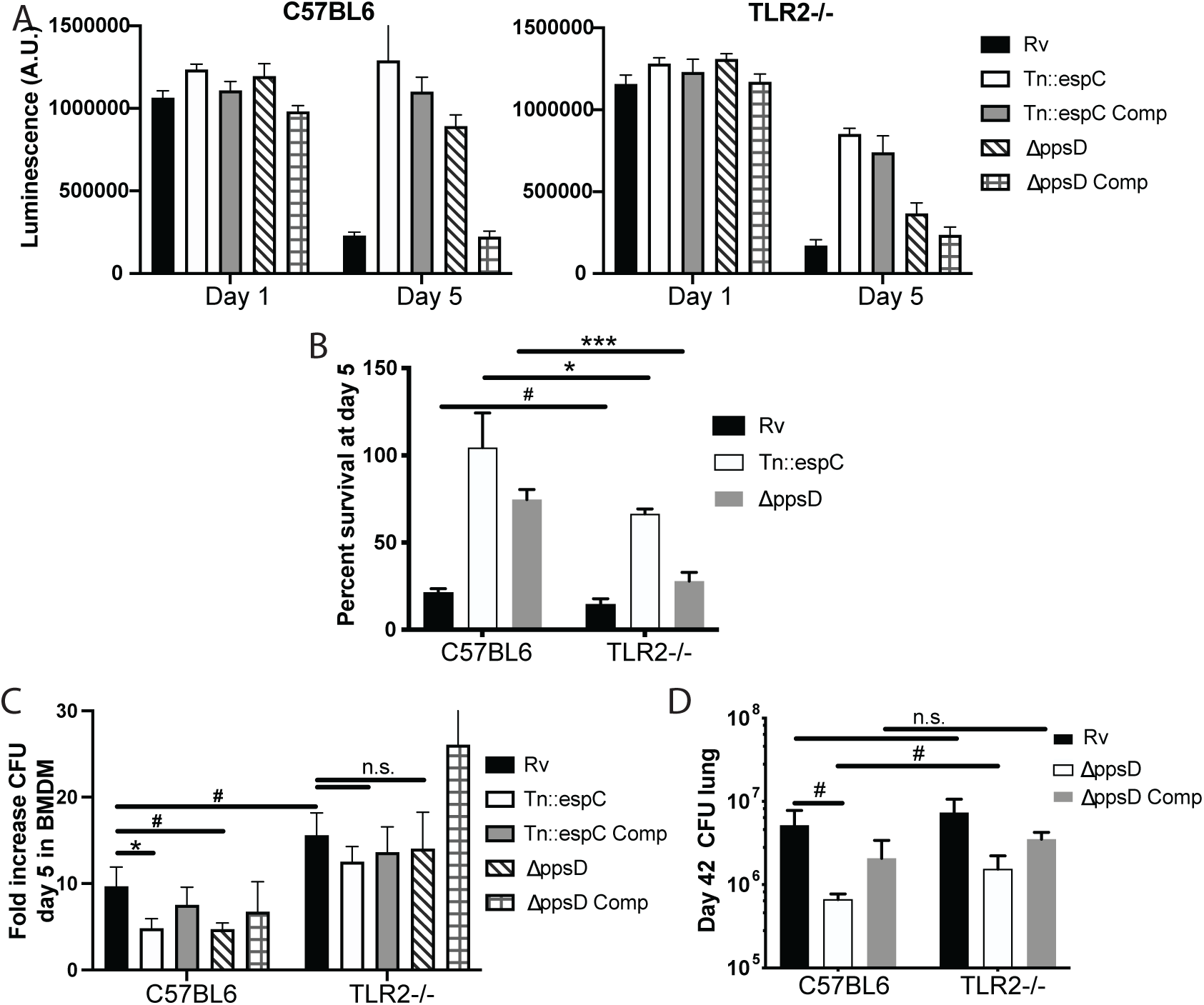
PDIM and ESX modulate TLR2-dependent infection outcomes in macrophages and mice. (A-C) The indicated BMDM were infected with the indicated Mtb strains at an MOI of 5:1 (A-B) or 2:1 (C). (A-B) Cell survival was determined using a CellTiterGlo luminescence assay at the indicated days post-infection. (C) At day 5 post-infection, cells were washed, lysed, and plated for CFU. (A-C) Mean +/-SD for 4 replicates. #p-value < 0.05, *p-value<0.01 unpaired two-tailed t-test. (D) C57BL/6J or TLR2-/-mice were infected with ∼200cfu of the indicated Mtb strains. 42 days post-infection, mice were euthanized and lungs were harvested and plated in serial dilutions to determine CFU Mean +/-SD for 5 mice per condition (one C57BL6/ppsD plate discarded for mold contamination-4 replicates for that condition). #p-value < 0.05, unpaired two-tailed t-test. (A-C) one of three independent experiments (D) one of two independent experiments.

We next sought to test whether the interaction between PDIM/ESX-1 and TLR2 impacts Mtb survival and growth in macrophages. PDIM and ESX-1 mutants have an attenuated growth phenotype in macrophages [30, 31]. We hypothesized that if TLR2-dependent responses contribute to this growth restriction, PDIM and ESX-1 mutants should grow more robustly in TLR2-/-BMDM than in wild-type BMDM. Alternatively, if PDIM and ESX-1 mutant growth restriction is entirely independent of TLR2-dependent responses, those mutants should grow similarly in wild-type and TLR2-/-BMDM. Using a low MOI to minimize induction of macrophage cell death, we infected wild-type or TLR2-/-BMDM with our wild-type or mutant Mtb strains. As expected, PDIM and ESX-1 mutants grew less well than wild-type Mtb in C57BL/6J BMDM (**Fig. 7C**). Growth of wild-type Mtb was modestly enhanced in TLR2-/-macrophages relative to wild-type macrophages; further, the PDIM and ESX-1 mutants grew significantly more robustly in the TLR2-/-BMDM than in wild-type BMDM, with growth similar to wild-type Mtb (**Fig. 7C**). These results suggest that TLR2-dependent responses contribute to growth restriction of the PDIM and ESX-1 mutants in macrophages. Together with our macrophage survival data, these results indicate that PDIM and ESX-1-mediated interference with TLR2-dependent responses contributes to the pathogenesis of Mtb infection in macrophages.

### PDIM and ESX-1 modulate TLR2-dependent infection outcomes in mice

Several studies have assessed the role of TLR2 in infection outcomes in mice. Some studies have shown a modest increase in Mtb growth in TLR2-/-mice relative to wild-type mice [59-61], while others have shown no difference in CFU. Most studies have shown increased lung pathology in TLR2 knockout mice, with larger infiltrates, less organization, and more inflammatory cells described as key features in various studies [59, 61, 62]. TLR2-/-mice have been shown to have increased Mtb dissemination to liver and spleen [59] and a more rapid progression to death following infection [59, 62, 63]. In aggregate, these data suggest that TLR2 plays a somewhat modest role in host control of TB infection. We hypothesized that PDIM and ESX-1 interference with the late TLR2-dependent response limits the contribution TLR2 makes to host control of Mtb infection. To test this hypothesis, we compared infection outcomes in wild-type and TLR2-/-mice. As previous profiling had demonstrated that CFU and pathology diverge by 6 weeks post-infection [61], we selected this timepoint for study. A limited number of mutant mice could be obtained for these studies; we thus focused on the interaction between PDIM and TLR2.

C57BL/6J and TLR2-/-mice were infected with wild-type, PDIM-mutant, and complemented PDIM-mutant Mtb (**Supp. Fig. S7A**). At the 6-week timepoint, as expected, PDIM-mutant Mtb growth was restricted relative to wild-type Mtb growth in C57BL/6J mice. Wild-type Mtb grew similarly in the lungs of C57BL/6J and TLR2-/-mice. In contrast, PDIM-mutant Mtb had increased growth in the lungs of TLR2-/-mice relative to C57BL/6J mice (**Fig. 7D**), suggesting that PDIM-mediated interference with activation of components of the TLR2-dependent response contributes to the capacity of the bacterium to grow in lung. Histopathologically, the lungs of C57BL/6J mice infected with wild-type Mtb demonstrated defined areas of inflammation by 6 weeks post-infection (**Supp. Fig. S7B**), with dense inflammatory infiltrates composed of foamy and non-foamy macrophages and lymphocytes clusters (**Supp. Fig. S7C**). Lungs of C57BL/6J mice infected with PDIM-mutant Mtb showed trends toward fewer areas of inflammation and smaller lesion sizes (**Supp. Fig. S7B)**. Examination of the regions of cellular infiltration in mice infected with PDIM-mutant Mtb were notable for similar presence of foamy macrophages and lymphocytes (**Supp. Fig. S7C**). In TLR2-/-mutant mice infected with PDIM-mutant Mtb, infiltrates showed a trend toward more numerous and larger areas of involvement than was observed in C57BL/6J mice (**Supp. Fig. S7B-C**). In total, our data suggest that PDIM modulation of TLR2-dependent responses contributes to pathogenesis *in vivo*. Growth of the PDIM mutant is only partially restored in TLR2-/-mice, indicating that additional mechanisms contribute to the attenuation of PDIM mutants *in vivo*.

## DISCUSSION

Accumulating data suggest that pathogenic bacteria evolve strategies for evading the components of immunity most critical for controlling their survival and replication. Multiple intracellular pathogens damage the phagosomal membrane in the course of pathogenesis, raising the question of whether this shared function reflects a convergent evolutionary strategy. While phagosomal membrane damage has been proposed to benefit the bacterium in the host-pathogen standoff, the mechanisms through which that benefit might accrue have not been well-established experimentally. In the case of pathogens recognized by TLR2, our results raise the possibility that damaging the phagosomal membrane may serve as a common strategy to limit effective inflammation.

Previous work has suggested that TLR2 can signal from the plasma membrane or endosome. Investigation of the mechanisms and consequences of TLR2 signaling has primarily focused on pathogen-specific induction of type I IFNs [7-10] and TNF [6, 64], largely in response to *S. aureus* exposure or viral infection. Our results support a model in which TLR2 activation in fact drives distinct compartment-specific pro-inflammatory transcriptional responses, reflected in both the sets of genes expressed and the kinetics of induction. Although a fundamental feature of TLR2 signaling, the two transcriptional response components are likely to have different relevance in the context of individual infections. In the case of TB, TNF, a component of the early response, is known to be critical for infection control. However, the later response component includes expression of multiple genes demonstrated to be important for both cell intrinsic control of Mtb and priming of the adaptive immune response. Our results demonstrating an interaction between TLR2 and PDIM or ESX-1 for infection outcome suggest that undermining this second component contributes to Mtb’s success as a pathogen. More broadly, our results suggest that expanding beyond TNF and type I IFNs as markers of TLR activation may offer both new insights into mechanisms and consequences of TLR signaling and into the links between TLR activation and control of pathogenic infection.

Our work suggests one potential mechanism through which PDIM and ESX-1 contribute to the pathogenicity of Mtb. PDIM has previously been shown to interfere with an effective MYD88 inflammatory response, as measured by the outcomes of macrophage recruitment to *M. marinum*-containing lesions *in vivo* and iNOS production [65]. In that work, this effect was hypothesized to be attributable to the unmasking of TLR agonists on the mycobacterial surface in the absence of PDIM, an abundant outer membrane lipid. Our results confirm an effect of PDIM on MYD88-dependent inflammation but point toward a different potential molecular interaction between PDIM and inflammation-namely that PDIM limits an endosomal component of the TLR2 response by limiting phagosome acidification. Two lines of evidence support the latter proposed mechanism. First, the effect we observe on TLR2-dependent inflammation is shared between PDIM and ESX-1. ESX-1-mediated secretion is not known to be required for localization of any known TLR2 agonist and in fact Mtb has multiple distinct TLR2 ligands; thus the common effect of PDIM and ESX-1 on TLR2-dependent inflammation is unlikely to be due to masking of TLR2 agonists. Second, PDIM and ESX-1 only minimally impact expression of the early TLR2-dependent gene cluster, suggesting that the inherent capacity of TLR2 to “recognize” cognate ligand on the bacterium is similar between wild-type Mtb and ESX-1 or PDIM mutant Mtb.

Together with previous work identifying Mtb factors that interfere with TLR2 activation, our work points toward an explanation for the puzzling disparity between the number of identified TLR2 ligands that Mtb possesses and the relatively modest phenotype of Mtb infection in TLR2 knockout mice. Work from other groups has identified mycobacterial strategies for interfering with TLR2 activation, primarily studied through an impact on TNF expression and release. The secreted hydrolase Hip1 has been shown to blunt the secretion of cytokines, including TNF-α, following infection [66]. Recently, the surface lipid sulfolipid-1 was shown to interfere with surface recognition of TLR2 agonists [67]. Our work suggests that the canonical Mtb virulence factors PDIM and ESX-1 function to blunt a distinct, endosome-specific component of the TLR2 response, and that this interference in fact modulates infection outcomes in macrophages and in mice. In aggregate, these results suggest that mycobacteria have evolved multiple strategies to undermine TLR2 activation from both the surface and endosome. Adding to the complexity of TLR2-dependent phenotypes in TB infection, studies of the relationship between Mtb TLR2 agonists and IFN-g-dependent functions have shown that TLR2 activation can dampen IFN-g-dependent gene expression and cell functions [38, 68]. Ultimately developing new strategies for treating tuberculosis will rely on a deep understanding of the pathogenesis of infection that enables the rational selection of therapeutic targets. Host-directed therapies enhancing the host-protective components of TLR2 activation might offer a path to a more effective inflammatory response to Mtb and ultimately more effective sterilization of TB infection.

## Materials and Methods

### Bacterial strains and culture

The indicated *Mtb* strains were grown in Middlebrook 7H9 broth (Difco) with Middlebrook OADC (BD), 0.2% glycerol, and 0.05% Tween-80. Mtb strains H37Rv, Tn::eccCa1, ΔppsD, ΔppsD::pMV261:: ΔppsD, and Δmas were characterized in Barczak et al [12]. Tn::espC was grown from a published transposon library in H37Rv [12]. The complement was generated by cloning the espACD operon from H37Rv into the Kpn and XbaI sites of shuttle plasmid pMV261 [69] (F primer atgacagatcggcctagctagg R primer attgtgagcccagtcgggaaa).

### Macrophage infections

BMDM were isolated and differentiated in DMEM containing 20% FBS (Cytiva) and 25ng/ml rm-M-CSF (R&D Systems) as previously described [12]. Infections were carried out as previously described [12, 70]. Briefly, *Mtb* strains used were grown to mid-log phase, washed in PBS, resuspended in PBS, and subjected to a low-speed spin to pellet clumps. BMDM were infected at the indicated MOI, allowing 3-4 hours for phagocytosis. Cells were then washed once with PBS, and media was added back to washed, infected cells. The MOI used for each time point was selected to maximize signal while minimizing infection-associated cell death.

### Mouse strains

C57BL/6J (Jackson Laboratories strain #000664), BALB/c (Jackson Laboratories strain #000651), *Sting*^-/-^ (C57BL/6J-*Sting*^*gt*^/J, Jackson Laboratories strain # 017537), *TLR4*^*-/-*^ (Jackson Laboratories strain #007227) *Tlr2*^*-*/-^ (B6.129-*tlr2*^*tm1Kir*^/J, Jackson Laboratories strain #004650), *MyD88*^-/-^ (B6.129P2(SJL)-*Myd88*^*tm1*.*1Defr*^/J, Jackson Laboratories strain # 009088), *TNFAR-/-* (B6.129S-*Tnfrsf1a*^*tm1Imx*^ *Tnfrsf1b*^*tm1Imx*^/J, Jackson Laboratories strain #003243), and *Trif-/-* (C57BL/6j-*Ticam1*^*Lps2*^/J, Jackson Laboratories strain #005037) mice were used for the preparation of bone marrow-derived macrophages.

### RNA isolation and qPCR

Infected BMDM were lysed at designated time points following infection with β-ME-supplemented Buffer RLT (Qiagen). RNA was isolated from lysate using an RNEasy kit (Qiagen) supplemented with RNase-free DNase I digest (Qiagen), both according to manufacturer’s protocol. cDNA was prepared using SuperScript III (Thermo Fisher Scientific) according to manufacturer’s protocol. qPCR was performed using PowerUP SYBR Green (Thermo Fisher Scientific) and primers specific to investigated genes relative to GAPDH control.

### RNA-Seq

Poly(A) containing mRNA was isolated from 1μg total RNA using NEBNext Poly(A) mRNA Magnetic Isolation Module (New England Biolabs). cDNA libraries were constructed using NEBNext Ultra II Directional RNA Library Prep Kit for Illumina and NEBNext Multiplex Oligos for Illumina, Index Primers Sets 3 and 4 (New England Biolabs). Libraries were sequenced on an Ilumina NextSeq500. Bioinformatic analysis was performed using the open source software GenePattern [59, 63]. Raw reads were aligned to mouse genome using TopHat, and Cufflinks was used to estimate the transcript abundance. FPKM values obtained by Cufflinks were used to plot heatmaps [71]. Three biological replicates for each condition were performed; replicates that failed QC metrics were not included in heatmaps and clustering. K-means clustering was performed in R and functional analysis was performed using IPA [18] (QIAGEN Inc., https://www.qiagenbio-informatics.com/products/ingenuity-pathway-analysis). RNAseq data is accessible on the NCBI GEO website GSE144330.

### TNF ELISAs

BMDM were infected at an MOI of 5:1 as described above. Following a 4 hour phagocytosis, BMDM were washed, and BMDM media was added back. At 24 hours post-infection, supernatants were collected for quantitation of TNF-α using an ELISA Ready-SET-Go! kit according to the manufacturer’s protocols. (ThermoFisher Scientific). Four replicates were performed per condition based on the determination that this would give 80% power to detect a 20% difference between samples.

### Imaging of pHrodo-labeled bacteria in macrophages

Imaging of pHrodo-labeled Mtb as described in Queval et al. [50]. Mtb strains were grown to mid-log phase, then washed twice with an equal volume of PBS. Mtb was then resuspended in 100mM NaHC03 with 0.5M pHrodo dye (Invitrogen) and incubated at room temperature in the dark for 1 hour. The labeled cells were then washed three times with PBS, after which BMDM infections were performed as described above. At the indicated time post-infection, infected BMDM were washed with PBS and fixed in 4% paraformaldehyde. Nuclei were labeled with DAPI (1.25μg/ml). Cells were then imaged on a Zeiss Elyra microscope with a 40x oil objective or a TissueFAXS confocal microscope with a 40x objective. Images were imported into CellProfiler [51] for analysis. Bacterial outlines were identified based on GFP signal; the outline was then expanded by 5 pixels, and pHrodo fluorescence intensity within the expanded outline was determined.

### CFU quantitation

Bacteria were prepared as described above, and added to BMDM at an MOI of 2:1. After 4 hours, cells were lysed in 0.5% triton X-100, diluted in 7H9 media, and plated on 7H10 plates for colony enumeration. For gentamicin killing assay, cells were treated with dynasore (80μM) prior to infection where indicated. Following the 4 hour phagocytosis, cells were washed in PBS with gentamicin (32μg/ml), then resuspended in bone marrow macrophage media with gentamicin (32μg/ml). After allowing 2 hours for killing of extracellular bacteria, cells were washed, lysed, diluted, and plated for colony enumeration.

### Quantitation of MFI for pHrodo-labeled zymosan beads

C57BL/6J BMDM were plated in an 8-chamber slide. Cells were pre-treated with concanamycin A, (50μM) or DMSO carrier for 15 minutes. Media was then removed, and pHrodo red zymosan bioparticles (Invitrogen) were added at 0.5mg/ml in BMDM media with concanamycin A or DMSO carrier. After 2 hours, media was removed and cells were washed once with PBS. Cells were then fixed in 4% paraformaldehyde and stained with DAPI (1.25μg/ml). Cells were imaged on a Zeiss Elyra PS.1 microscope with a 20x objective. Images were analyzed using a CellProfiler image analysis pipeline. DAPI-stained nuclei were identified and counted and integrated red pHrodo fluorescence was measured for each image.

### Macrophage survival assays

C57BL/6J or TLR2-/-BMDM were plated in 96-well format and infected with Mtb strains at an MOI of 5:1. Cells were harvested day 1 and day 5 post-infection using a CellTiter-Glo Luminescent Cell Viability Assay Kit (Promega) in accordance with the manufacturer’s instructions. After gentle agitation and 10-minute incubation, wells were read on a Tecan Spark 10M luminescent plate reader. Media was replenished every 48 hours post-infection.

### Mouse infections

C57BL/6J (Jackson Laboratories strain #000664) or TLR2-/-(Jackson Laboratories strain # 021302) mice were infected via low dose aerosol exposure with an AeroMP (Biaera Technologies). 3-5 mice per condition were harvested at day 0 to quantify inoculum. 6 weeks post-infection, mice were euthanized in accordance with AALAC guidelines, and lungs were harvested for CFU and histopathology. Formalin-fixed lungs were embedded in paraffin, sectioned, and stained with hematoxylin and eosin by the MGH histopathology core. Images were acquired on a TissueFAXS slide scanner (TissueGnostics).

## Acknowledgements

The authors would like to thank Drs. Roi Avraham, Bryan Bryson, Sarah Fortune, and Jonathan Kagan for critical manuscript review and Dr. Lenette Lu and the laboratories of Drs. Marcia Goldberg and Cammie Lesser for helpful discussions. We would additionally like to thank Dr.Sabine Ehrt for the pckA mutant, parent, and complement strains. The work was funded in part by an MGH Transformative Scholar Award (AKB) and made possible by help from the Harvard University Center for AIDS Research (CFAR), an NIH funded program (P30 AI060354).

## Author Contributions

Conceptualization, A.E.H. and A.K.B, Methodology, A.E.H., C.J., and A.K.B., Investigation, A.E.H., C.J., S.C.P., W.R.Z., and A.K.B, Analysis, N.J and A.J.M., Writing, A.E.H., C.J. and A.K.B, Funding Acquisition, A.K.B.

