## Supplementary Figures for "*Mycobacterium tuberculosis* canonical virulence factors interfere with a late component of the TLR2 response"

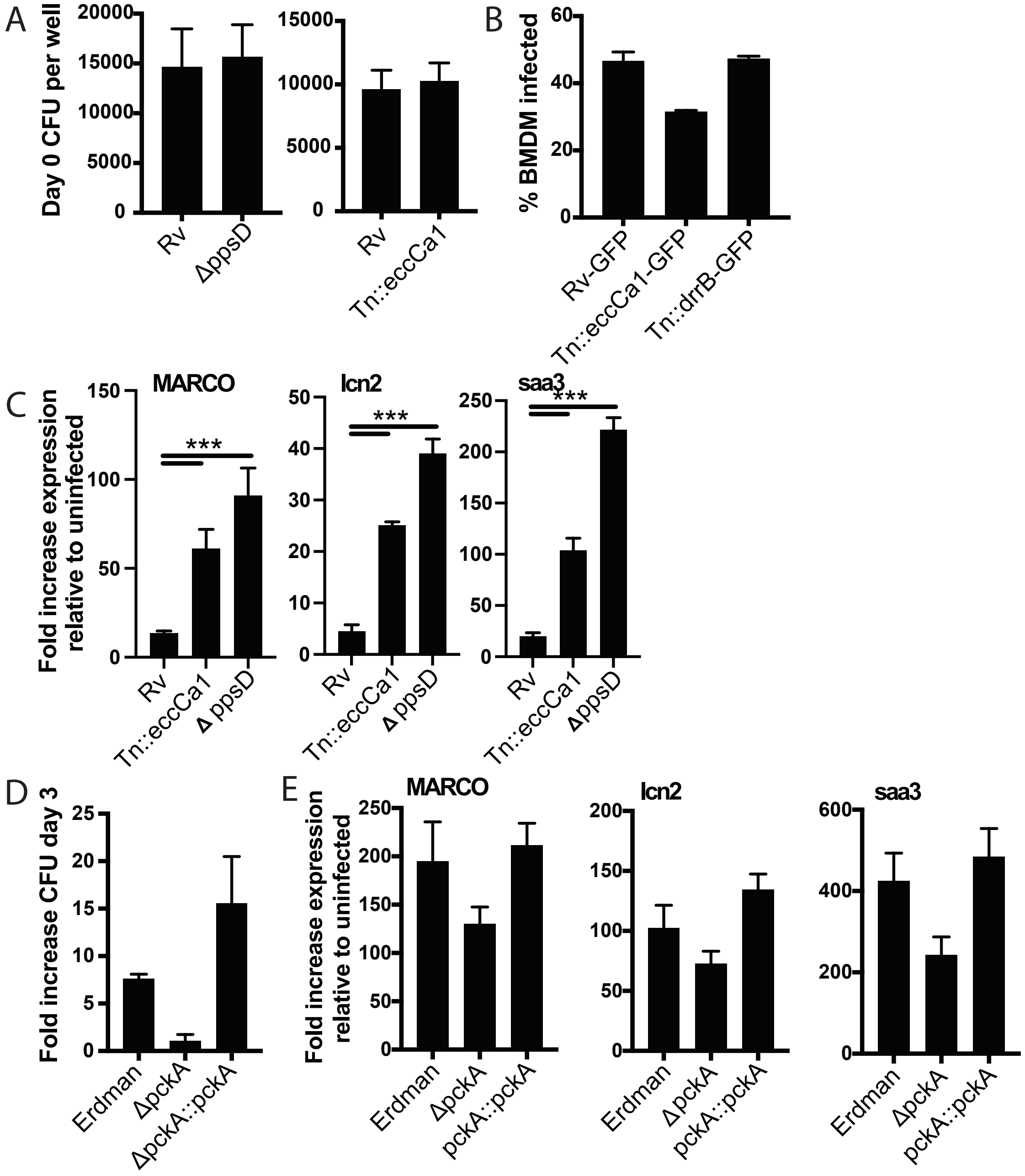


**Supp. Fig. S1.** (A) C57BL/6J BMDM were infected with Mtb at an MOI of 2:1. After 4 hours of phagocytosis, cells were washed, lysed, and plated for CFU. (B) C57BL/6J BMDM were infected with the indicated Mtb strains expressing GFP at an MOI of 5:1. After 4 hours of phagocytosis, BMDM were washed and analyzed by flow cytometry to quantitate the percent of cells with detectable Mtb-GFP signal. (C, E) BALB/c (C) or C57BL/6J (E) BMDM were infected with the indicated Mtb strains at an MOI of 2:1. RNA was harvested 24 hours post-infection, and expression of the indicated genes was quantified by qPCR relative to GAPDH control. (D) J774 cells were infected with the indicated Mtb strains at an MOI of 2:1. Cells were lysed at day 0 and day 3; lysates were plated for CFU. (E) C57BL/6J BMDM were infected with the indicated Mtb strains at an MOI of 2:1. (A, C-E) Mean +/- SD for 4 replicates. (B) Mean +/- SD for 3 replicates.

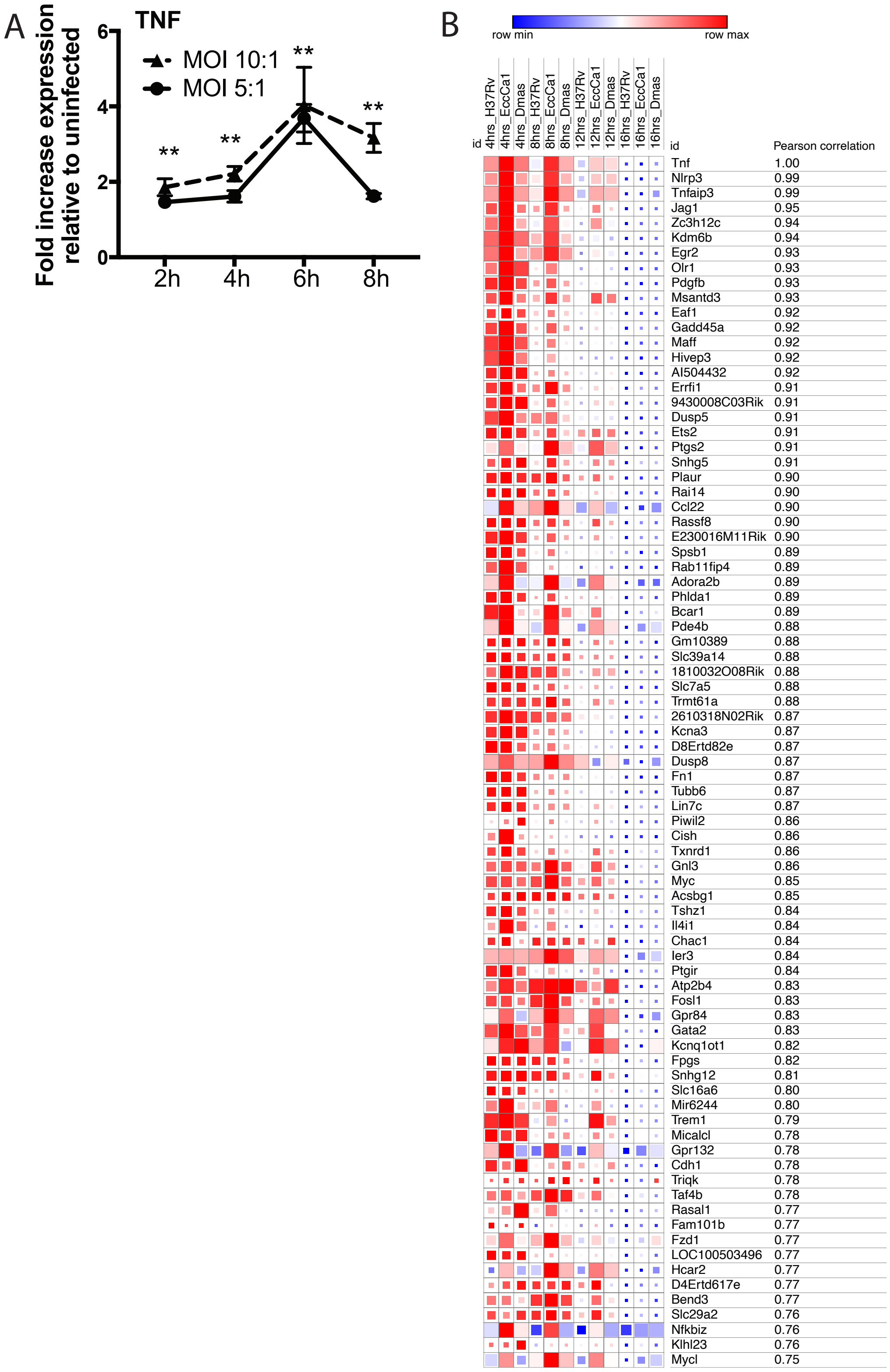


**Supp. Fig. S2.** (A) C57BL/6J BMDM were infected with H37Rv at the indicated MOI. RNA was harvested at the indicated timepoints. Expression of TNF relative to GAPDH control was profiled using qPCR. **p-value < 0.001 for the comparison of either MOI with uninfected at a given timepoint. (B) correlation analysis for genes co-expressing with TNF was performed on the RNAseq dataset. 81 genes with a pearson correlation co-efficient >= 0.75 were identified. Blue-red gradient reflects row minimum to row maximum expression; box size represents reflects absolute expression level.

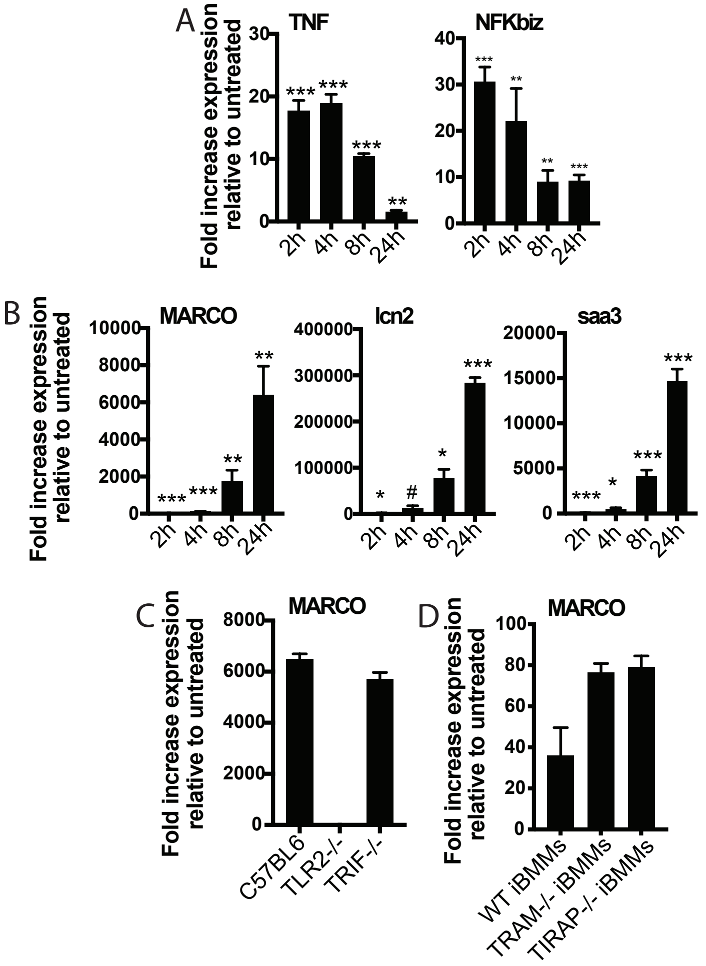


**Supp. Fig. S3.** (A-B) C57BL/6J BMDM were treated with PAM2CSK 1μg/ml and RNA was harvested at the indicated timepoints. (C) The indicated BMDM were treated with PAM3CSK 1μg/ml and RNA was harvested at 24 hours. (D) The indicated iBMMs were treated with PAM3CSK 1μg/ml and RNA was harvested at 24 hours. (A-D) qPCR was performed to quantify expression of the indiated genes relative to GAPDH control. Mean +/- SD for 4 replicates. #p-value < 0.05, *p-value < 0.01, **p-value < 0.001, ***p-value < 0.0001, unpaired two-tailed t-test for the comparison with untreated.

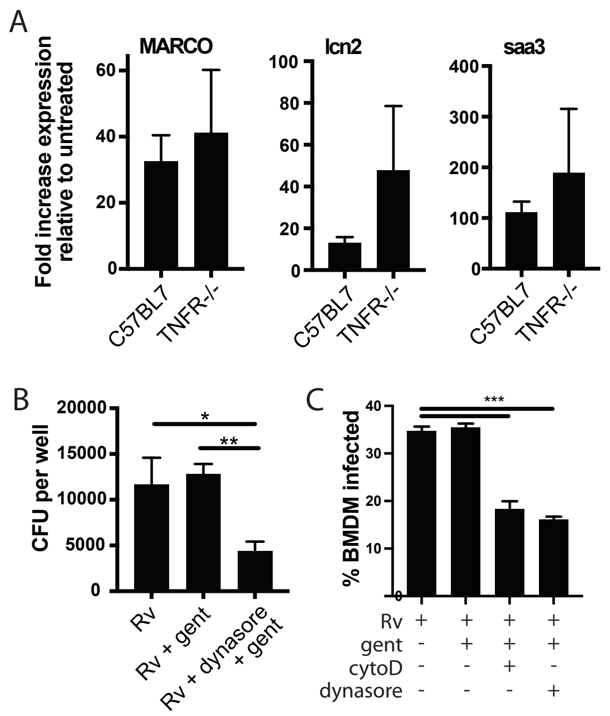


**Supp. Fig. S4.** (A) The indicated BMDM were infected with H37Rv at an MOI of 2:1. RNA was harvested 24h post-infection. Expression of the indicated genes was quantified by qPCR relative to GAPDH control. (B) C57BL/6J BMDM were pretreated with dynasore (80μM) where indicated for 30 minutes prior to infection. Cells were then infected with H37Rv at an MOI of 2:1. Following 4 hours of phagocytosis, cells were washed in PBS with gentamicin (32μg/ml) and resuspended in media with gentamicin (32μg/ml) for 2 hours. Cells were then washed with PBS, lysed, and plated for CFU. Mean +/- SD of 4 replicates. (C) C57BL/6J BMDM were pretreated with cytochalasin D (10μM), or dynasore (80μM) where indicated. Cells were then infected with Rv-GFP at an MOI of 5:1. Following 4 hours of phagocytosis, cells were washed in PBS and resuspended in media with gentamicin (32μg/ml) for 4 hours. Cells were again washed with PBS, fixed, and analyzed by flow cytometry. Gates for Mtb-GFP uptake were set based on an uninfected control. Mean +/- SD for 3 replicates. *p-value < 0.01, **p-value < 0.001, unpaired two-tailed t-test.

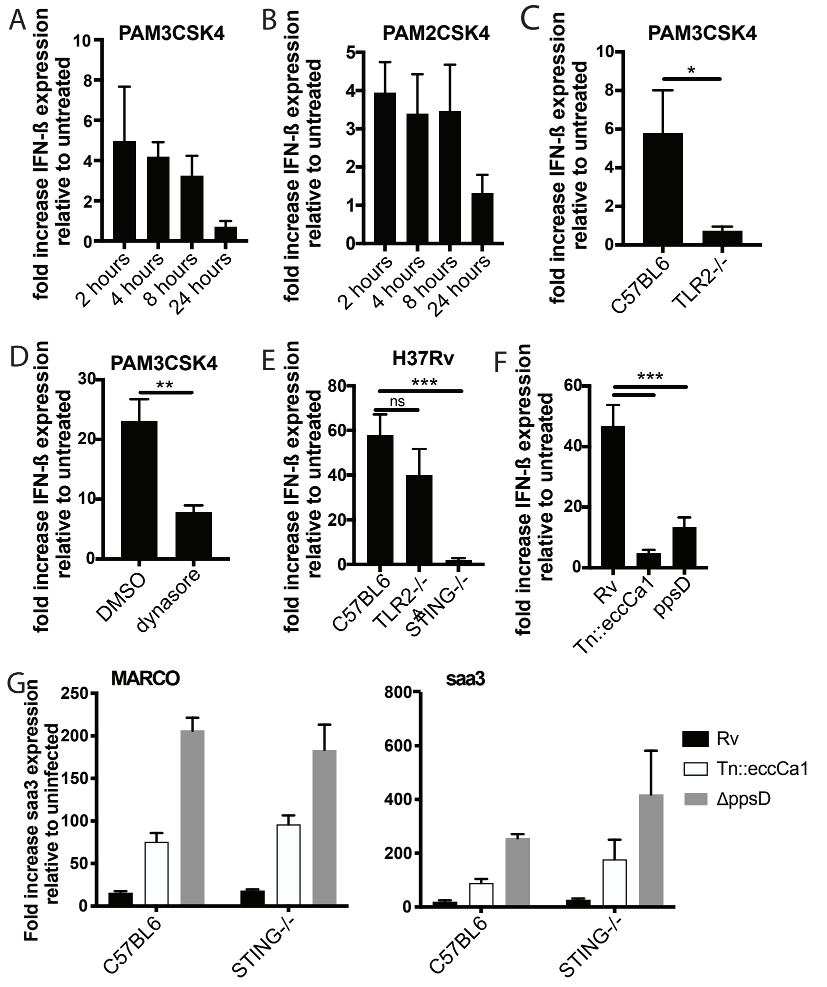


**Supp. Fig. S5.** (A-D) The indicated BMDM were treated with PAM2CSK4 (100ng/ml) as indicated. (D) Cells were pre-treated with dynasore (80μM) for 15 minutes prior to ligand treatment. (E-G) C57BL/6J yjr indicated BMDM (E, G) or C57BL/6J (F) were infected with the indicated Mtb strains at an MOI of 5:1 (E-F) or 2:1 (G). RNA was harvested at the indicated times (A-B), at 2 hours (C-D), 6 hours post-infection (E-F), or 24 hours post-infection (G). Expression of the indicated genes relative to GAPDH control was quantified by qPCR. Mean +/- SD for 4 replicates. *p-value < 0.01, **p-value < 0.001, ***p-value < 0.0001, unpaired two-tailed t-test.

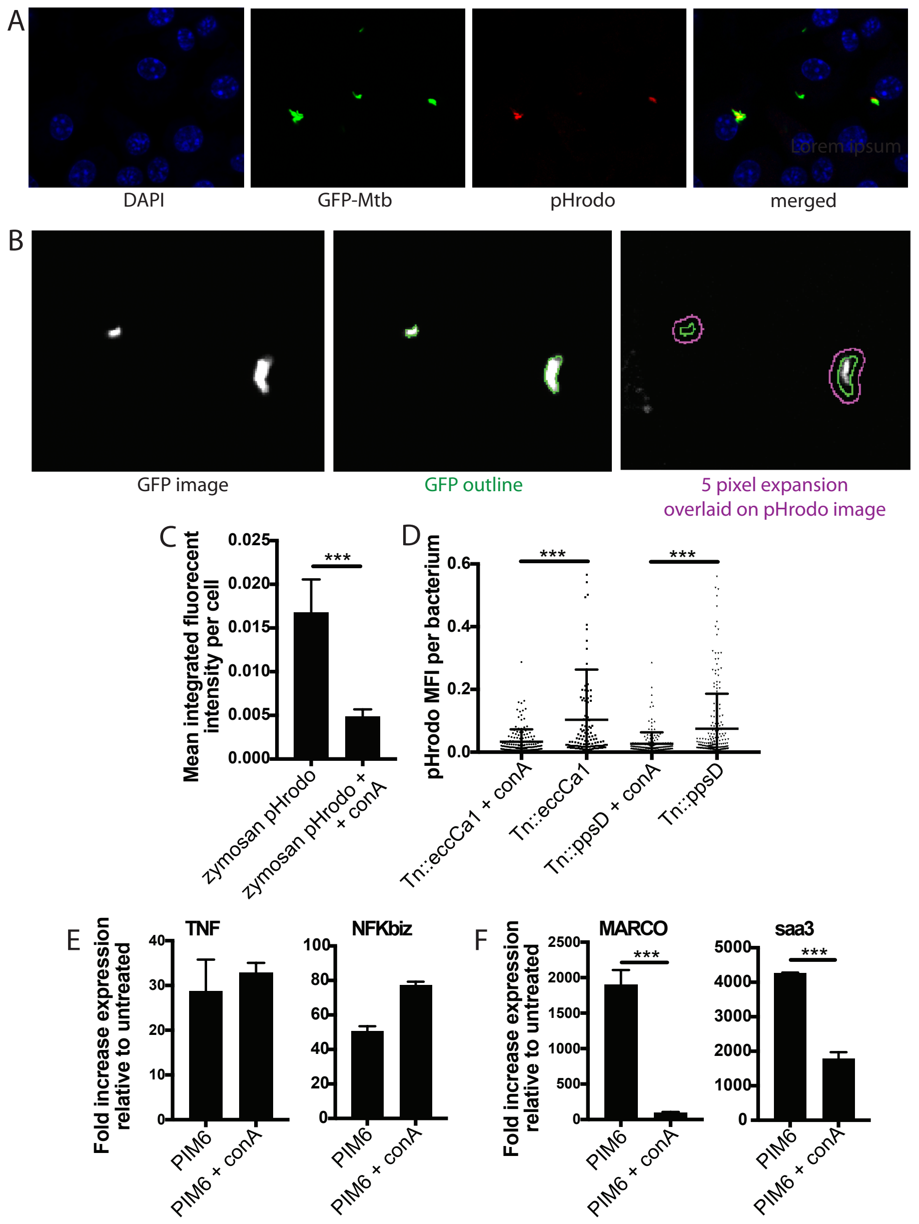


**Supp. Fig. S6.** (A-B) C57BL6 BMDM were infected with pHrodo labeled GFP-Mtb. (A) Example DAPI, GFP, and pHrodo images. (B) CellProfiler automated identification of GFP bacteria and expansion of 5 pixels around the GFP outline to capture all relevant pHrodo signal. (C-D) C57BL/6J BMDM were pre-treated with concanamycin A (50$\mu$M) or DMSO carrier as indicated. (C) pHrodo-labeled zymosan bioparticles were then added to the cells in BMDM media containing concanamycin A or carrier. Phagocytosis was allowed to proceed for 2 hours; cells were then fixed and imaged using a Zeiss Elyra PS.1 with a 20x objective. CellProfiler image analysis was used to quantitate integrated fluorescent intensity per cell. (D) The indicated Mtb strains expressing GFP were labeled with pHrodo and used to infect C57BL/6J BMDM at an MOI of 3:1. Cells were fixed 24 hours post-infection and imaged. Bacteria were identified based on GFP signal, and pHrodo mean fluorescence intensity was measured around each bacterium (as in Queval *et al,* 2017). A minimum of 128 bacteria (D) were analyzed per group. Mean +/- SD for 5 fields per condition. (E-F) C57BL/6J BMDM were pre-treated with concanamycin A (50$\mu$M) or carrier for 15 minutes, then treated with PIM6 (1$\mu$g/ml) in the presence of concanamycin A (50$\mu$M). RNA was harvested at 2 hours (E) or 24 hours (F) and expression of the indicated genes relative to GAPDH control was quantified by qPCR. Mean +/- SD of 4 replicates. *** p-value < 0.0001, unpaired two-tailed t-test.

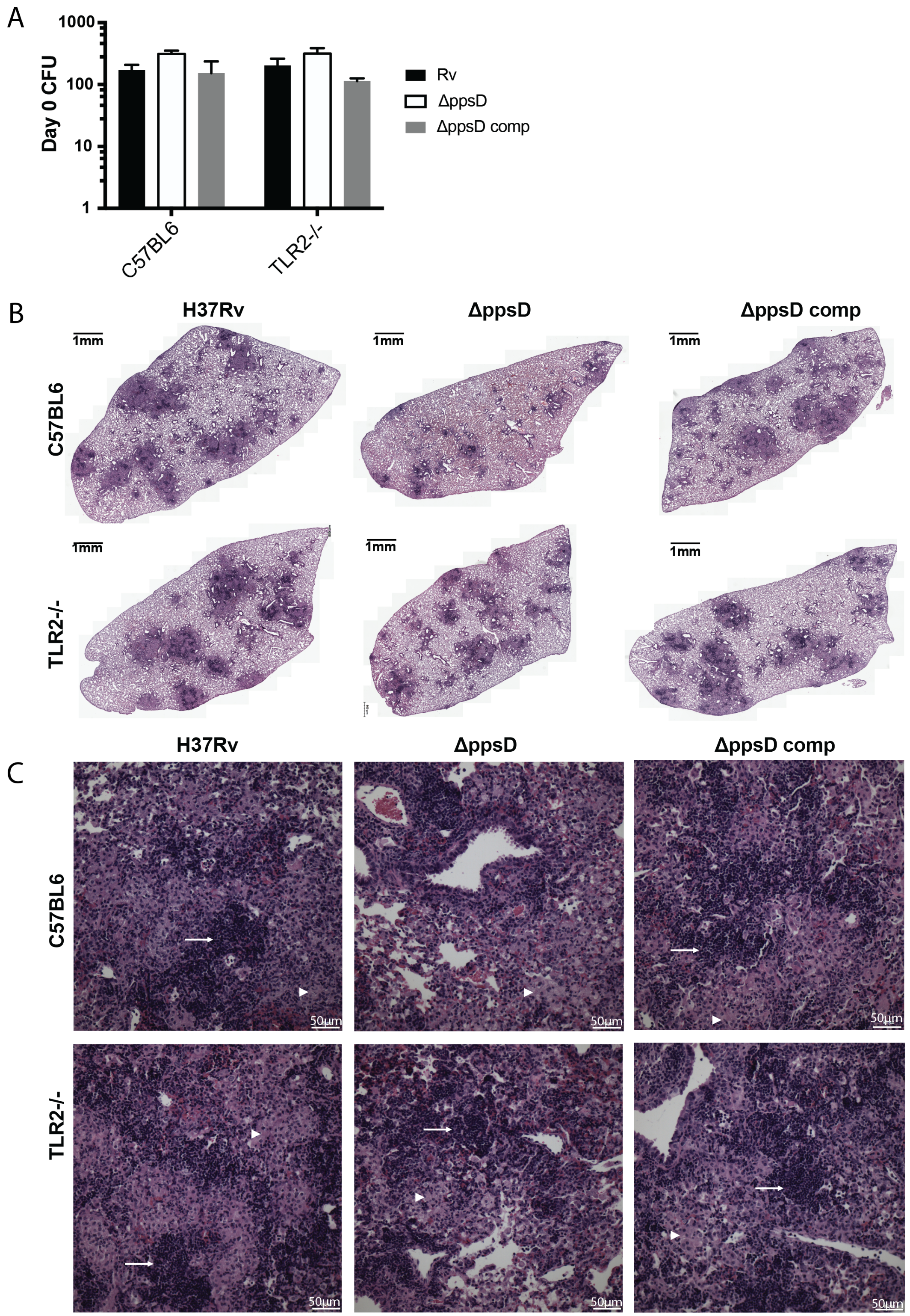


**Supp. Fig. S7.** The indicated mouse strains were infected with the indicated Mtb strains via low-dose aerosol infection. (A) Lungs were harvested after infection and plated on day 0 for CFU. Mean +/- SD for 3-5 mice per condition. (B-C) Lungs were fixed in formalin, embedded in paraffin, and sectioned. Sections were stained by hematoxylin & eosin and imaged on a TissueFACS Slide Scanner using a 20x objective. (B) Overview of all fields for a sectioned lung. (C) Individual fields of view for areas of cellular infiltration in each lung. Arrows: Areas of dense inflammatory cell infiltration. Arrow heads: areas of foamy macrophages. #p-value < 0.05, unpaired two-tailed t-test.
